# Modulating Sphingosine-1-Phosphate receptors to improve chemotherapy delivery to Ewing sarcoma

**DOI:** 10.1101/805655

**Authors:** Enrica Marmonti, Hannah Savage, Aiqian Zhang, Claudia Alvarez, Miriam Morrell, Keri Schadler

## Abstract

Tumor vasculature is innately dysfunctional. Poorly functional tumor vessels inefficiently deliver chemotherapy to tumor cells; vessel hyper-permeability promotes chemotherapy delivery primarily to a tumor’s periphery. Here we identify a method for enhancing chemotherapy delivery and efficacy in Ewing sarcoma (ES) in mice by modulating tumor vessel permeability. Vessel permeability is partially controlled by the G protein-coupled Sphinosine-1-phosphate receptors 1 and 2 (S1PR1 and S1PR2) on endothelial cells. S1PR1 promotes endothelial cell junction integrity while S1PR2 destabilizes it. We hypothesize that an imbalance of S1PR1:S1PR2 is partially responsible for the dysfunctional vascular phenotype characteristic of ES and that by altering the balance in favor of S1PR1, ES vessel hyper-permeability can be reversed. In this study, we demonstrate that pharmacologic activation of S1PR1 by SEW2871 or inhibition of S1PR2 by JTE-013 caused more organized, mature, and functional tumor vessels. Importantly, S1PR1 activation or S1PR2 inhibition improved chemotherapy delivery to the tumor and anti-tumor efficacy. Our data suggests that pharmacologic targeting of S1PR1 and S1PR2 may be a useful adjuvant to standard chemotherapy for ES patients.

**NOVELTY AND IMPACT:** This study demonstrates that Sphingosine-1-Phosphate (S1P) receptors are potential novel targets for tumor vasculature remodeling and adjuvant therapy for the treatment of Ewing Sarcoma. Unlike receptor tyrosine kinases that have already been extensively evaluated for use as vascular normalizing agents in oncology, S1P receptors are G protein-coupled receptors, which have not been well studied in tumor endothelium. Pharmacologic activators and inhibitors of S1P receptors are currently in clinical trials for treatment of auto-immune and cardiovascular diseases, indicating potential for clinical translation of this work.

## INTRODUCTION

Ewing sarcoma (ES) is an aggressive sarcoma of bone and soft tissue that represents 3% of all pediatric malignancies^1^. ES tumor vessels, like vessels in many solid tumors, are characterized by a discontinuous endothelial lining with wide endothelial cell junctions and disjointed pericyte coverage. This contributes to marked vessel leakiness^2^. Excessive vascular leakage causes chemotherapy extravasation at the periphery of the tumor, contributing to suboptimal chemotherapy efficacy^2^.

Vascular remodeling, or normalization, using anti-angiogenic agents enhances perfusion and drug delivery in solid tumors^3^. A “normalized” vascular phenotype includes improved vessel function and reduced hyper-permeability; specifically, increased perivascular cell coverage and restored endothelial barrier integrity^3^. It was recently demonstrated that combination of anti-angiogenic agent celecoxib with standard chemotherapy improved survival in patients with metastatic ES^4^. Although this study indicates the potential of vascular remodeling as adjuvant therapy for ES, severe toxicities limited clinical usefulness of the anti-angiogenic agent.

Here, we present an alternative approach to improve drug delivery by targeting tumor vascular hyper-permeability. Several tumor secreted factors such as vascular endothelial growth factor-A (VEGF-A) cause disruption of adherens junctions by phosphorylation, internalization and cleavage of Vascular-Endothelial cadherin (VE-cadherin), an essential regulator of endothelial cell-cell adhesive properties^5^. Sphingosine-1-Phosphate (S1P) is a sphingolipid that regulates endothelial barrier function and angiogenesis via G protein-coupled S1P Receptors 1 and 2 (S1PR1 and S1PR2)^6^. S1PR1-G_i_-Rac signaling decreases vessel permeability and enhances endothelial cell-to-cell junctions by inducing VE-cadherin trafficking to adhesion sites^6^. S1PR1 also regulates the interaction of perivascular and endothelial cells in microvasculature, creating more mature vessels^7^. Conversely, S1PR2 activation disrupts endothelial cell junctions via the G_12/13_-Rho-Rho kinase kinase (ROCK)-PTEN pathway by preventing VE-cadherin translocation to cell contact sites^8^. The balance of S1PR1 and S1PR2 in a vascular bed thus defines endothelial barrier integrity^8^. In physiologic conditions, a high S1PR1: low S1PR2 ratio maintains a stabilized and intact vascular endothelium while a low S1PR1: high S1PR2 ratio causes endothelial barrier dysfunction^9^. Although the antagonistic relationship between S1PR1 and S1PR2 is known in healthy endothelium and several disorders related to pathologic vascular permeability^8^, the role of these receptors in modulating tumor vasculature function is poorly understood.

We previously demonstrated that S1PR1 and S1PR2 are expressed on ES endothelium and that reduced tumor vessel hyper-permeability after exercise-induced shear stress correlates with increased S1PR1 and decreased S1PR2^10^. These findings suggest that ES vessel function might be improved by altering the balance in favor of S1PR1. To elucidate the role of S1PR1 and S1PR2 on tumor vasculature function, we performed preclinical studies in human ES mouse xenograft models using selective pharmacological modulators of S1PR1 and S1PR2. SEW2871 is a S1PR1-selective agonist that induces receptor internalization and subsequent recycling via AKT/ERK1/2/Rac1 pathway activation^6^. JTE-013 is an S1PR2 antagonist which prevent ROCK-PTEN pathway activation^8^.

Here, we demonstrate that activation of S1PR1 signaling or inhibition of S1PR2 signaling induced more normalized tumor vessels with reduced hyper-permeability. Importantly, vascular normalization by S1P receptor modulation correlated with significantly improved chemotherapy efficacy. This is the first study to demonstrate the role of S1P receptors in vascular function in ES. Our findings reveal a novel mechanism of tumor vascular remodeling via activation of S1PR1 and inhibition of S1PR2 signaling that contributes to increased chemotherapy delivery and, therefore, increased anti-tumor effects in mice.

## MATERIAL AND METHODS

### Cell culture

A673 ES cells (ATCC Cat# CRL-1598, RRID:CVCL_0080) were cultured per manufacturer recommendation. Cells are authenticated by STR analysis at MD Anderson Cancer Center Characterized Cell Line Core Facility within the lasts three years and routinely tested negative for mycoplasma contamination.

### Animals and experimental protocol

The Institutional Animal Care and Use Committee at The University of Texas MD Anderson Cancer Center approved the animal studies. A673 cells (2.5×10^6^) were injected subcutaneously into the backs of 6-week-old athymic nude (nu/nu) male mice. After tumors reached ∼50 mm^3^, mice were randomized into 4 groups: daily oral vehicle (dimethyl sulfoxide [DMSO] or alcohol), daily oral pharmaceutical agent (SEW2871 10mg/kg or JTE-013 1.5mg/kg; Cayman Chemical), intravenous doxorubicin (2mg/kg twice per week; Premier Pharmacy), and combination therapy with pharmaceutical agent + doxorubicin. The SEW2871 experiment was repeated in an orthotopic model using the same treatment schedule in which A673 cells (2.5×10^5^) were injected into the gastrocnemius of 6-week-old nude mice. Mice were housed in individually ventilated cages under pathogen-free conditions. Animals had free access to food and water and were kept on a 12-hour light/12-hour dark cycle.

### Vessel structure and function

Five minutes prior to euthanasia, tumor-bearing mice were injected via tail vein with 100μL tomato-lectin (2mg/mL in PBS 7.4, VectorLab) or high molecular weight FITC-Dextran (2,000,000 mol weight, 10mg/mL in PBS pH 7.4; Sigma-Aldrich).

Frozen tumor sections were stained with the following primary antibodies: rat anti-CD31 (1:50, BD Pharmingen), rabbit anti-alpha-smooth muscle actin (α-SMA, 1:100, Abcam Ab5694;), rabbit anti-NG2 Chondroitin Sulfate Proteoglycan (NG2, 1:100, AB5320 Millipore), mouse rat anti-Vascular Endothelial cadherin (Ve-caderin, 1:100, BD Pharmingen). Nuclei were stained with Fluoro-Gel II with DAPI (Electron Microscopy Sciences). Images were captured with a Leica DM5500 B upright microscope imaging system (Leica Microsystems) and analyzed using NIS-Elements Imaging Software. For all immunostaining assays, 5 random fields from each tumor sample were quantified as previously described^11^. Images for VE-cadherin and CD31 staining were obtained with a 63x oil immersion objective using a Zeiss LSM 880 with Airyscan FAST confocal microscope. Pearson’s correlation coefficient was calculated to measure the colocalization correlation of the intensity distribution between VE-cadherin and CD31 of 5 random vessels per tumor sections.

### Hypoxia

qPCR was performed with iQ SYBR^®^ Green Supermix (Bio-Rad) and run on a LightCycler^®^ 480 Instrument II (Roche). Carbonic anhydrase IX (CAIX) (FW: GAGAAGGCAGCACAGAAG G and REV: GGCTTCTCACATTCTCCAAGAT) and Glucose Transport-1 primers (GLUT-1) (FW: GGGCCA AGAGTGTGCTAAA and REV: CTTCTTCTCCCGCATCATCTG), Vascular Endothelial Growth Factor-A (VEGF-A) (FW: GTGAATGCAGACCAAAGAAAGATA G and REV: CCAGGACTTATACCGGGATTTC); Hypoxia inducible factor 1 subunit alpha-1α (HIF-1α) (FW: GTCTGCAACATGGAAGGTATTG and REV: GCAGGTCATAGGTGGTTTCT) as well as internal control primers Glyceraldehyde-3-phosphate dehydrogenase (GAPDH) (FW: AACAGCAACTCCCACTCTTC and REV: CCTGTTGCTGTAGCCGTATT) were synthesized by Integrated DNA Technologies.

### Proliferation assay (Live-cell imaging)

A673 cells (5×10^3^ per well) were grown in a 96-well plate and filmed every four hours for 48 hours with Incucyte live cell imaging system (Essen Instrument., Ann Harbor, MI). Doxorubicin (0.01nM), SEW2871 (50nM), or JTE-013 (50μM) was added at the beginning of the quantification period, or DMSO or ethanol as a negative control. The experiment was performed three times. Proliferation was monitored by analyzing the cell occupied area (% confluence) of images over time.

### Statistical analysis

All values are reported as means ± standard error of the mean. Statistical significance of the results was calculated by two-way analysis of variance. Intergroup differences were evaluated by using student’s t-test and linear mixed models. The statistical analysis was performed using SpSS (version 21) and Graphpad software.

All data and detailed methods will be made available upon reasonable request.

## RESULTS

### S1PR1 activation by SEW2871 promotes tumor vascular normalization

To determine the effect of S1PR1 activation in ES, A673 subcutaneous and orthotopic xenografts were established in nude mice. Mice were treated with an S1PR1 selective agonist, SEW2871, alone or in combination with doxorubicin. There were no significant differences in microvessel density or total vessel count between tumors in different treatment groups in either subcutaneous or orthotopic tumors (data not shown). In both models, S1PR1 activation promoted tumor vessel normalization. SEW2871 treatment stimulated the formation of more elongated vessels with a greater number of open lumens in subcutaneous tumors (Elongated Vessels: SEW p=0.016, Lumens: SEW p=0.046, Fig. 1A and C) and orthotopic tumors (Elongated Vessels: SEW p=0.014, Lumens: SEW p=0.013, Fig. 1B and D). Further, S1PR1 activation significantly increased mural cell coverage of tumor vessels. The percentage of tumor capillaries with observable alpha-smooth-muscle actin (α-SMA) (SEW p=0.002, Fig. 1E) and Neuron-glial 2 (NG2) labeling (SEW p=0.046, Fig.1F) was significantly increased in SEW-treated subcutaneous tumor sections compared to untreated tumors. Similarly, but with a less significant effect, α-SMA (SEW p=0.061, Fig. 1E) and NG2 (Fig. 1F, SEW p=0.065) coverage of capillaries was also increased in SEW-treated orthotopic tumors.

**Figure 1.**
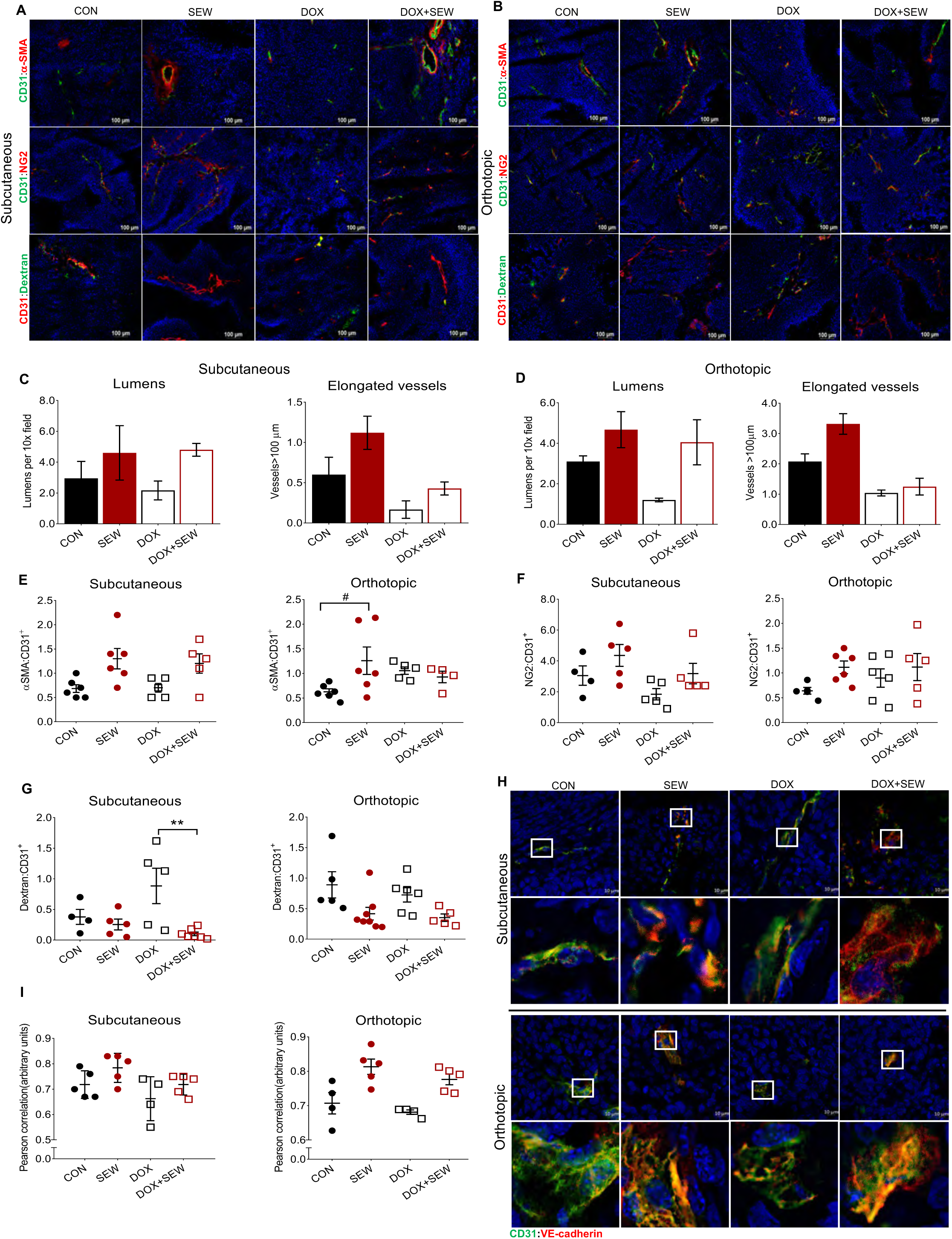
S1PR1 activation by SEW2871 promotes tumor vascular normalization. After A673 tumor cells injection (7 days post-subcutaneous injection and 11 days post-intramuscular injection), tumor-bearing mice were treated with doxorubicin [DOX] (2mg/Kg, twice per week, intravenously) and/or SEW2871 [SEW] (S1PR1 agonist, 10mg/Kg, daily, orally). **(A and B)** α-SMA (red) or NG2 (red) and CD31 (green) or FITC-dextran leak (green) and CD31 (red) immunofluorescence with DAPI staining (nuclei) in subcutaneous **(A)** and orthotopic tumors **(B)**; scale bar: 100μm. **(C and D)** The number of visible lumens and the number of vessels>100μm (large) were counted in 5 random sections/tumor. **C.** *Subcutaneous.* Bars show means ± SEM, *n=*5-6. Two-way ANOVA, open lumens (SEW p=0.0465), vessels>100μm (SEW p=0.016, DOX p=0.001). **D.** *Orthotopic.* Bars show means ± SEM, *n=*5-7. Two-way ANOVA, open lumens (SEW p=0.014), vessels>100μm (SEW p=0.015, DOX p<0.0001). **(E)** Mean αSMA:CD31 ratio ± SEM calculated in 5 random sections/tumor. *Subcutaneous*, two-way ANOVA (SEW p=0.002), n=5-6. *Orthotopic, t*wo-way ANOVA (SEWxDOX p=0.047), *n=*5-6. Post-hoc Tukey test and indicated by #p=0.061. **(F)** Mean NG2:CD31 ratio ± SEM calculated in 5 random sections/tumor. *Subcutaneous*, two-way ANOVA (SEW p=0.046, DOX p=0.068), n=4-5. *Orthotopic, t*wo-way ANOVA (SEW p=0.065), *n=*5-6. **(G)** Mean Dextran:CD31 ± SEM ratio for individual A673 tumors. *Subcutaneous, t*wo-way ANOVA (SEW p=0.009, DOXxSEW p=0.050), n=4-7. Post-hoc Tukey test and indicated by **p<0.01, ***p<0.001. *Orthotopic*, two-way ANOVA (SEW p=0.004), n=5-8. **(H)** Representative confocal images of subcutaneous and orthotopic tumors showing colocalization of VE-cadherin (red) and CD31 (green). Nuclei were couterstained with DAPI (blue). Scale bar 10μm. Part of the section (white rectangle) is shown below in higher magnification (300x). **(I)** Graph shows Pearson’s correlation coefficient for VE-cadherin and CD31 markers in subcutaneous and orthotopic tumors represented in Figure 1H. For each tumor 5 vessels were analyzed in different optical regions. *Subcutaneous*, two-way ANOVA (DOXO p=0.044, SEW p=0.044). *Orthotopic*, two-way ANOVA (SEW p=0.0003).

Tumor vascular permeability was assessed by injection of high molecular weight FITC-dextran (2000 KDa), a molecule that does not leak from functional vessels^12^. S1PR1 activation caused a 34% reduction in vascular leakiness of subcutaneous tumor vessels that became more evident in doxorubicin-treated mice (87%; DOXOxSEW p=0.005, Fig. 1G). In the orthotopic model, a significant reduction in dextran leakage was observed in both tumors treated with SEW2871 alone (53%) or in combination with doxorubicin (52%; SEW p=0.004, Fig. 1G).

To confirm improvement in barrier integrity by the S1PR1 activation, we analyzed VE-cadherin protein expression and localization at the plasma membrane by confocal microscopy analysis. Although SEW2871 treatment did not change the total VE-cadherin expression levels (data not shown), it did significantly increase the translocation of VE-cadherin at the adhesion sites, demonstrated by co-localization with CD31 in both subcutaneous and orthotopic tumors (Fig. 1H-I).

### S1PR2 inhibition by JTE-013 promotes tumor vasculature normalization

To study the impact of S1PR2 inhibition on tumor angiogenesis, mice bearing subcutaneous A673 tumors were treated with JTE-013, doxorubicin, or the combination of JTE-013 and doxorubicin. Inhibition of S1PR2 caused a significantly higher microvessel density (p=0.027) that appeared more elongated (p=0.068) with a greater number of open lumens (p=0.067) in subcutaneous tumors (Fig. 2A, C). There was no significant effect of JTE-013 in remodeling microvessel structure and organization in orthotopic tumors (data not shown). Although α-SMA (Fig. 2A, D) and NG2 cell coverage did not change (Fig. 2A, E), JTE-013 reduced tumor vessel leakage by 68% and 43.7% in subcutaneous and orthotopic tumors, respectively (p=0.038, Fig. 2F and 2G). Additionally, while the pharmacological inhibition of the S1PR2 pathway did not affect VE-cadherin translocation at the plasma membrane (data not shown), we observed a trend toward S1PR2 antagonism increasing VE-cadherin protein expression levels in both subcutaneous (SEW p=0.064, Fig. 2H-I) and orthotopic tumors (SEW p=0.071, Fig. 2H-I).

**Figure 2.**
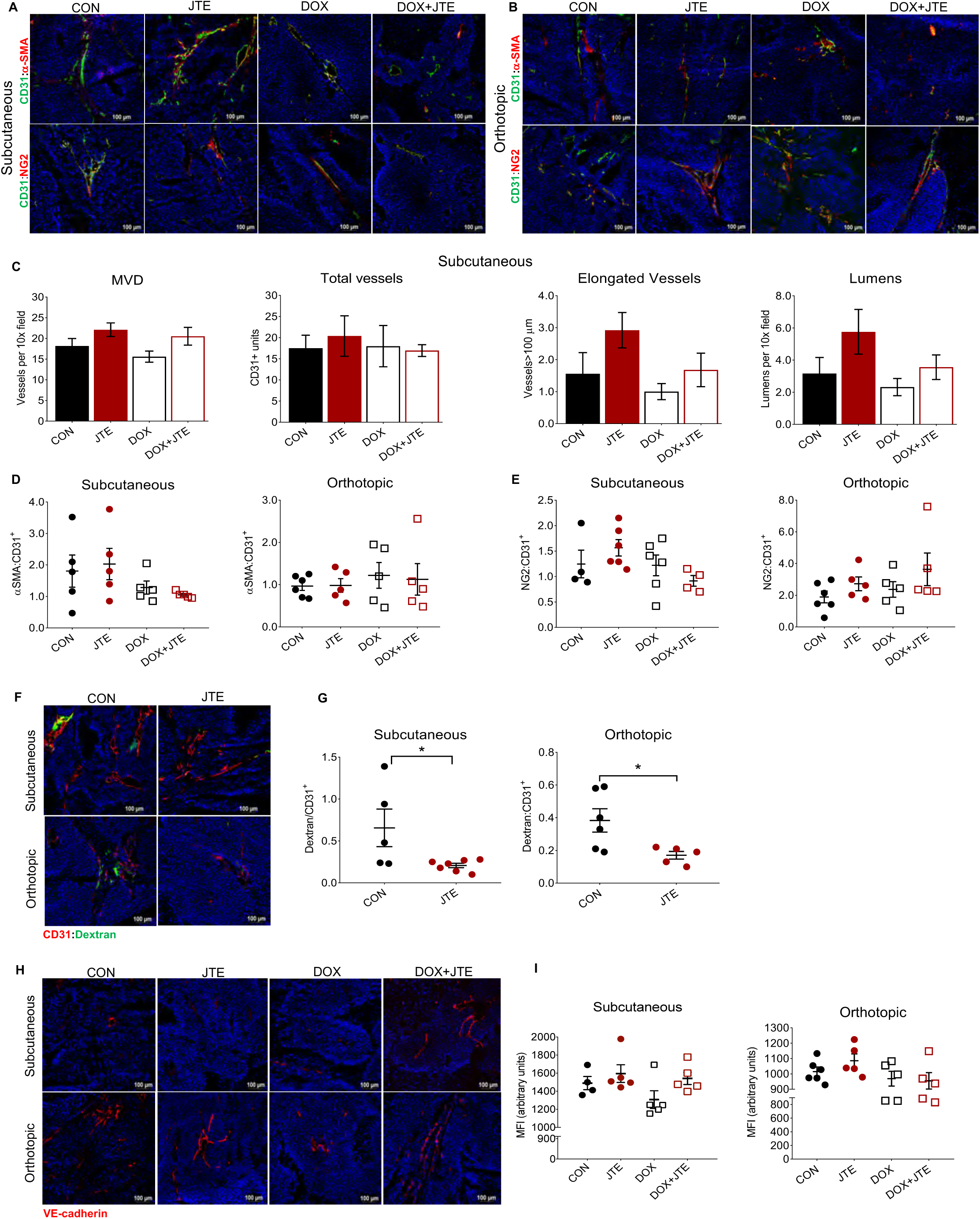
S1PR2 receptor inhibition by JTE-013 promotes tumor vasculature normalization. After A673 tumor cells injection (7 days post-subcutaneous injection and 13 days post-intramuscular injection), tumor-bearing mice were treated with doxorubicin [DOX] (2 mg/Kg, twice per week, i.v.) and/or JTE-013 [JTE] (S1PR2 antagonist, 2.5mg/Kg daily in subcutaneous model and 5mg/Kg twice a day in orthotopic model, orally). **(A, B)** α-SMA (red) or NG2 (red) and CD31 (green) or VE-cadherin in subcutaneous **(A)** and orthotopic tumors **(B)**; scale bar: 100μm. **(C)** The average number of microvessels density (MVD), total vessels, the number of visible lumens and the number of vessels>100μm (large) were counted in 5 random sections/subcutaneous tumor. Bars show means ± SEM, *n=*5-6. Two-way ANOVA, MV (JTE p=0.023); total vessels (p=non-significant); open lumens (JTE p=0.067); vessels>100μm (JTE p=0.068, DOX p=0.103). **(D)** Mean αSMA:CD31 ratio ± SEM calculated in 5 random sections/tumor. *Subcutaneous*, two-way ANOVA (DOXO p=0.061), n=5. *Orthotopic, t*wo-way ANOVA (p=ns), *n=*5-6. **(E)** Mean NG2:CD31 ratio ± SEM calculated in 5 random sections/tumor. *Subcutaneous*, two-way ANOVA (DOXO p=0.109), n=4-6. *Orthotopic, t*wo-way ANOVA (JTE p=0.107), *n=*5-6**. (F)** Representative images of FITC-dextran leak (green) and CD31 (red) immunofluorescence with DAPI staining (nuclei) in orthotopic and subcutaneous tumors; scale bar: 100μm. **(G)** Mean Dextran:CD31 ± SEM ratio for individual A673 tumors. *Subcutaneous, n=*5-7, T-test Student *p=0.038. *Orthotopic*, T-test Student *p=0.027, n=5-6. **(H)** Representative images of subcutaneous and orthotopic tumors. VE-cadherin (red), DAPI staining (nuclei); scale bar: 100μm. **(I)** Quantification of VE-cadherin by mean fluorescence intensity (MFI) calculated in 5 random sections/tumor. Values represent means ± SEM. *Subcutaneous*, two-way ANOVA (JTE p=0.071), n=5-6. *Orthotopic*, two-way ANOVA (JTE p=0.064), n=5-6.

### S1PR1 activation or S1PR2 inhibition improved chemotherapy efficacy

The vascular remodeling observed after SEW2871 treatment correlated with a 2-fold increase (51%) in the number of functional (lectin perfused) vessels compared to control tumors (p=0.028, Fig. 3A). Consistent with improved vascular function leading to reduced tumor hypoxia, SEW2871 significantly reduced CaIX (46%; SEW p=0.043, Fig. 3B) and VEGF-A mRNA levels (71%; SEW p=0.045) and had a clear trend toward reduced GLUT-1 (63%, Fig. 3B) and HIF1-α mRNA in orthotopic tumors (33%, Fig. 3B). Improved tumor perfusion and reduced hypoxia should improve chemotherapy delivery. Indeed, SEW2871 plus chemotherapy inhibited A673 tumor growth better than chemotherapy alone (subcutaneous: 41% better, Time: DOX+SEW vs DOX p<0.0001; and orthotopic: 45% better, Time: DOX+SEW vs DOX p<0.0001, Fig. 3C). No additive effect of SEW2871 and doxorubicin in inhibiting proliferation of A673 cells was observed *in vitro* (Fig. 3D).

**Figure 3.**
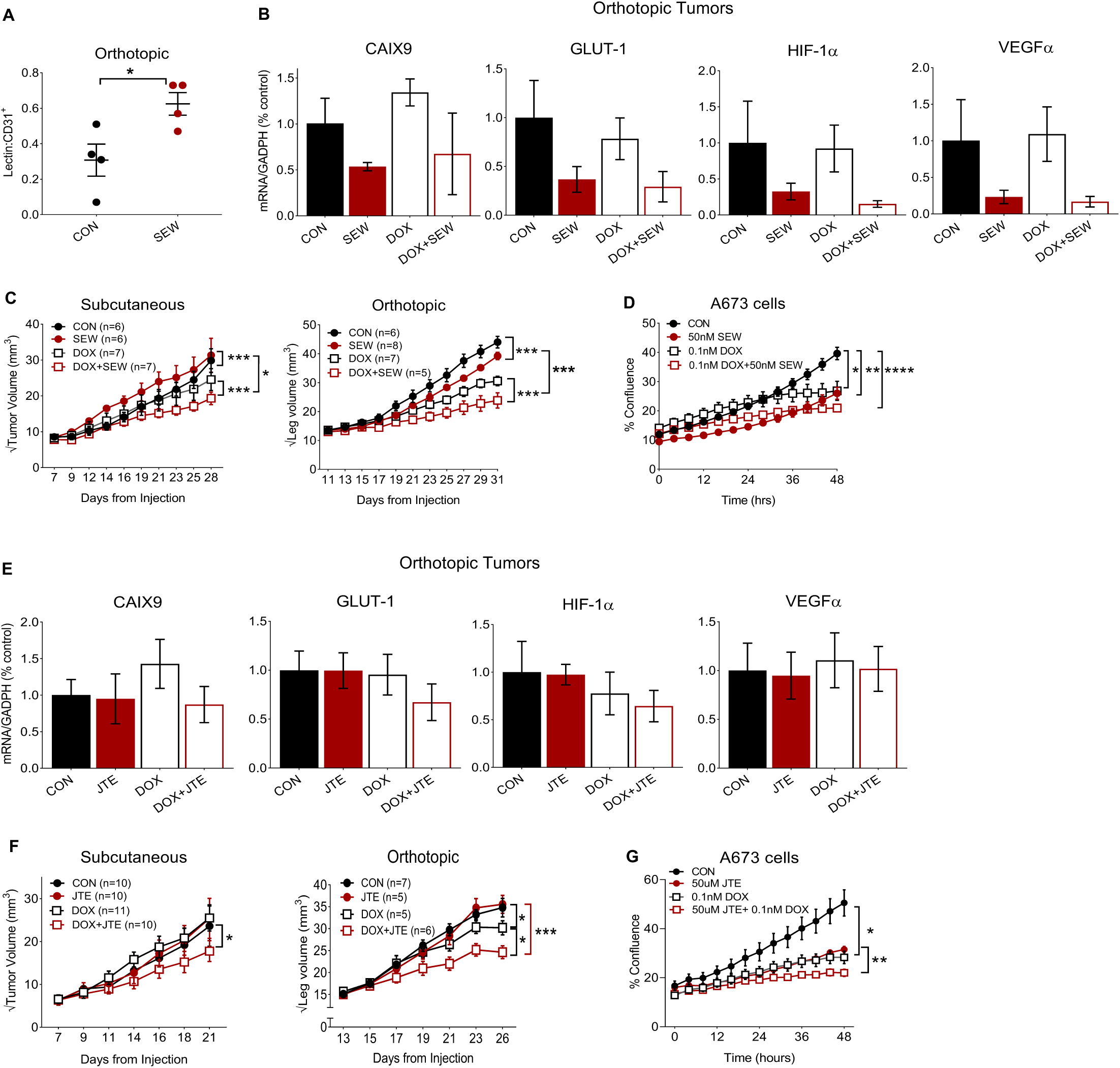
S1PR1 activation by SEW2871 or S1PR2 inhibition by JTE-013 improves chemotherapy efficacy. **(A)** Mean Tomato-lectin:CD31 ± SEM (n=4) ratio defined the percentage of perfused vessels, t-test *p=0.028. **(B)** RT-PCR analysis of the mRNA expression levels of CaIX, GLUT-1, HIF1-α, VEGF-A normalized against GADPH in orthotopic tumor homogenates. Data are expressed as the mean ± SEM of triplicate values. Bars represent means ± SEM, *n =*5-6. Statistical significance of the results was calculated by two-way analysis of variance (CaIX: SEW p=0.043; GLUT-1: SEW p=0.0511; VEGF-A: SEW p=0.045; HIF-1α: SEW p=0.078). **(C)** A673 tumor volumes from mice treated with [DOX] and/or [SEW] were measured in indicated days. *Subcutaneous.* Values are means ± SEM for 6-7 animals in each group. Linear Mixed Model (Time: SEW vs CON ***p<0.0001; Time: DOX+SEW vs CON \**p* = 0.0131 and Time: DOX+SEW vs DOX \*\*\**p =* <0.0001). *A673 Orthotopic.* Values are means ± SEM for 5-8 animals in each group. Linear Mixed Model (TIME: CON vs DOX *p<0.05; Time: CON vs SEW **p<0.01; Time: DOX+SEW vs CON \*\*\**p* <0.0001). **(D)** A673 cell proliferation assay (expressed as % of cell confluence) treated with SEW2871 (50nM) and doxorubicin (0.1nM) and combination of the two drugs. Linear Mixed Model: CON vs SEW ***p<0.001; CON vs DOXO ***p<0.001. **(E)** RT-PCR analysis of the mRNA expression levels of CaIX, GLUT-, HIF-1α, VEGF-A normalized against GADPH in orthotopic tumor homogenates. Data are expressed as the mean ± SEM of triplicate values. Bars represent means ± SEM, *n=*5-6. Statistical significance of the results was calculated by two-way ANOVA (CaIX: p=ns; GLUT-1: p=ns; VEGF-A p=ns; HIF-1α p=ns). **(F)** A673 tumor volumes from mice treated with [DOX] and/or [JTE] were measured in indicated days. *Subcutaneous.* Values are means ± SEM for 6-8 animals in each group. Linear Mixed Model: Time: DOXO+JTE vs JTE *p=0.020. *Orthotopic.* Values are means ± SEM for 5-7 animals in each group. Linear Mixed Model: CON vs DOX p=0.016; DOX vs DOXO+JTE p=0.032. **(G)** A673 cell proliferation assay (expressed as % of cell confluence) when treated with JTE-031 (50μM) and DOX (0.1nM) and combination of the two drugs. Linear Mixed Model: CON vs DOX *p<0.05; JTE vs DOX+JTE **p<0.01.

Unlike S1PR1 activation, S1PR2 pathway inhibition did not significantly change CaIX, GLUT-1, HIF1-α, VEGF-A mRNA levels in orthotopic tumors (Fig. 3E). Still, tumors treated with the combination of JTE-013 and doxorubicin were significantly smaller than tumors treated with doxorubicin alone in both xenograft models (subcutaneous: 43.9%, DOX+JTE vs JTE p=0.02; orthotopic 49.8% better, p=0.03) (Fig. 3F). JTE-013 treatment did not enhance the cytotoxicity of doxorubicin in A673 cells *in vitro* (Fig. 3G).

## DISCUSSION

Here, we provide evidence that modulating the ratio of S1PR1:S1PR2, by inhibiting S1PR1 or activating S1PR2, may be a novel method to remodel tumor vasculature and increase the delivery and thus the efficacy of chemotherapy.

In ES tumors, the activation of S1PR1 preferentially manifested as more elongated and open lumen vessels markedly covered with mural cells. Instead of promoting vascular sprouting and neoformation as previously reported in PyMT breast cancer^13^, the activation of S1PR1 by the agonist SEW2871 principally remodeled the morphology of ES tumor vessels to a more mature phenotype. This different vascular response may be partially dependent on tumor type. In comparison to other tumor types, ES vascular network is unique as it is characterized by a more morphologically organized vascular network^10^. However, consistent with other tumors, vasculature is dysfunctional and hyper-permeable.

The inhibition of S1PR2 by the antagonist JTE-013 only marginally increased the microvessel density, consistent with a previous report in S1PR2^-/-^ mice bearing Lewis Lung carcinoma and B16BL6^14^. Since more significant effects on vascular remodeling was observed by activating S1PR1, the inhibition of S1PR2 by pharmacological modulation might be not sufficient enough to overcome the vascular dysfunction largely caused by lack of S1PR1 pathway activations in ES tumors.

In addition, we demonstrated that improved endothelial cells junction function might be involved in tumor vascular normalization. Activation of S1PR1 reduced leakage of tumor vessels by stimulating the recruitment of mural cells around the endothelium and increasing VE-cadherin localization at endothelial junctions. Inhibition of S1PR2 also decreased vascular hyper-permeability, but the mechanism by which this occurs is less clear, as changes to VE-cadherin localization were subtle. The trend toward increased VE-cadherin expression after inhibition of S1PR2 is consistent with previous reports that S1PR2 signaling via NF-κB^15^ promotes the expression of the zinc-finger transcription factor Snail that represses VE-cadherin transcription in endothelial cells exposed to cancer cell-conditioned media^16,17^.

In addition, we demonstrated that the S1PR1 agonism by SEW2871 contributes to increased blood vessel perfusion and oxygenation of the tumor. Lower levels of hypoxia responsive transcripts, including VEGFα, confirm a reduction in hypoxia^18^. Considering that lower levels of VEGFα result in more stable endothelial cell adhesion junctions^5,19^, S1PR1 activation may alternatively increase vessel function as a result of decreased tumor hypoxia.

Moreover, since hypoxia induces cellular adaptations that promote cell survival and resistance to chemotherapy^20^, the reduced hypoxic environment likely contributes to a greater anti-tumor response. By using athymic nude mice, which lack functional T-cells, we removed any possible effects of S1P receptor signaling modulation on lymphocytes trafficking and immune regulation^21^. Therefore, the increased chemotherapy efficacy observed by the combined therapy was likely endothelial cell dependent.

Our findings demonstrate that S1PR1 and S1PR2 in ES vasculature can be modulated to normalize tumor vessels and improve chemotherapy efficacy. As such, the pharmacologic targeting of S1PR1 and S1PR2 warrants further study as potential adjuvant therapy targets for Ewing sarcoma patients.

## Abbreviations

S1P: Sphingosine-1-Phosphate
S1PR: Sphingosine-1-Phosphate receptor
ES: Ewing Sarcoma
VEGF-A: Vascular Endothelial Growth Factor-A
Akt: Protein kinase B
ERK: Extracellular signal-regulated kinase
Rac1: RAS-related C3 botulinum toxin substrate 1
VE-cadherin: Vascular Endothelial-cadherin
ROCK1: Rho associated protein kinase1
PTEN: Phosphatase and tensin homologue
α-SMA: alpha-smooth muscle actin
NG2: Neural/glial antigen2 Chondroitin Sulfate Proteoglycan
CAIX: Carbonic anhydrase IX
GLUT-1: Glucose Transport-1
HIF1α: Hypoxia inducible factor 1 subunit alpha
GADPH: Glyceraldehyde3-phosphate dehydrogenase
PyMT: Polyoma middle T antigen
DOXO: Doxorubicin
SEW: SEW2871
JTE: JTE-013
DMSO: dimethyl sulfoxide
DAPI: 4’,6’-diamidino-2-phenylindole, dyhydrochloride
NF-κB: Nuclear factor kappa light chain enhancer of activated B cells
FITC: Fluorescein isothiocyanate

## ACKNOWLEDGMENTS

We acknowledge Minjeong Park, for help with statistical analysis, the Center for Energy Balance at the M.D. Anderson Cancer Center, and funding from the Cancer Prevention Research Institute of Texas (CPRIT) grant number RP190256.

